# Distributions of threshold crossing times of messenger RNA

**DOI:** 10.64898/2026.08.20.745891

**Authors:** Arunendra Kumar Verma, Hillol Kumar Barman, Krishna Rijal, Dibyendu Das

**Affiliations:** Department of Physics, Indian Institute of Technology Bombay, Powai, Mumbai 400076, India; Department of Organismic and Evolutionary Biology, Harvard University, Cambridge, MA, USA

## Abstract

Within the studies of stochastic gene expression, apart from the variability of copy number of gene products, the problems of threshold crossing of those products are biologically important as they often lead to terminal cellular events. Here, we study the threshold crossing problem of the messenger ribonucleic acid (mRNA) and present an exact probability distribution of first passage times in Laplace space. The function furnishes moments of any order and also predicts the characteristic time of the exponential tail of the distribution, which we match against Gillespie simulations. We find that all the measures of relative fluctuations of the threshold crossing times show U-shapes within this simple model of mRNA, as was found earlier in more mathematically involved models of threshold crossing time statistics of proteins. Furthermore, we extend the exact formula to include the phenomenon of DNA duplication and the corresponding doubling of transcription rate. As expected, the distribution varies considerably depending on the onset of the duplication stage within the cell cycle.

## I. INTRODUCTION

Variability of gene products across cells is an active area of research in cell biology. It happens due to intrinsic causes like stochastic gene expression^1–6^, as well as extrinsic causes like cell partitioning error and division time uncertainties across generations^7–15^. With technologies of single-cell RNA sequencing and single-molecule fluorescence in situ hybridization to quantify messenger RNA (mRNA) levels, their stochastic variations in cells may be followed. Similarly, the variability of protein levels is studied using fluorescent proteomic imaging, mass cytometry, and mass spectrometry. Such experimental studies have revealed enormous intercellular variability within isogenic populations^16–22^. The variability has strong implications for medical problems, like cancer drug resistance and microbial persistence^23–27^. It has been shown how fluctuations in gene expression may persist across generations and may render reproductive fitness to drug-resistant populations independently of mutations, and that may be viewed theoretically as a stochastic first passage problem^28^. Apart from this, first passage arises in other gene product level crossing problems to be discussed below, and this forms the main focus of this paper.

In any stochastic process, the question of *first passage* arises when the process is terminated on reaching a boundary^29^. The statistics of the corresponding time before reaching the target is of great interest and is referred to as ‘first passage time’ (FPT). In biology there are many first passage problems which have been of interest – for example, the first encounter of remote parts of DNA^30^, protein search for sites on DNA^31^, promoter site exposure with nucleosome kinetics^32^, recovery after backtracking for proofreading during RNA synthesis^33^, motile kinetochore capture by multiple microtubules^34^, and other processes like molecular rapture, self-assembly of proteins attaining a critical size, and fixation or extinction events in species populations^35^.

Within the area of gene expression, which is of our interest in this paper, first passage problems of threshold crossing of gene products have been of interest as they often trigger crucial biological events. When *λ*-bacteriophage infects E. coli, and the holin protein expressed from *λ*-DNA crosses a threshold, causing the bacterial cell lysis, the event has been considered as a first passage – experimental lysis times have been compared with exact theoretical predictions^36–40^. While crossing the blood-brain-barrier or lung epithelial cells, the bacteria Streptococcus Pneumonia traps itself in endosomes which occasionally perforates when the toxin Pneumolycin secreted by the bacteria crosses a threshold – the experimental pore formation times have been matched with theoretical FPT distributions^41^. The process of cell division has been treated as a first passage problems with the size (or protein biomass) of cells crossing a threshold^42–45^.

The problem of threshold crossing of gene products, apart from computational investigations, has been studied in many works that yield FPT moments analytically, but the full and exact FPT distributions are known only in a few cases. For example, even for the basic protein synthesis through a two-stage model, with transcription at rate *k* of mRNA and its degradation at rate *γ*, and translation from mRNA to protein at rate *k*_*p*_ with its degradation at *γ*_*p*_^46^ – the FPT distribution is not known explicitly. For the geometrically distributed bursty protein production limit, often suitable for bacteria and yeast when *γ*_*p*_ ≪ *γ*^46–48^, various exact results have been recently found, which we highlight here. The moments of the FPT distribution were obtained in^37^ with arbitrary autoregulation of genes. Subsequently, the full FPT distribution of protein ‘number’ crossing a threshold was obtained^39^ in the absence of autoregulation – the results were applied to the pore formation times in endsomes as discussed above^41^. For protein ‘concentrations’ crossing a threshold appropriate for growing cells with changing volume, the suitable full FPT distribution for bursty protein production was derived and compared to experimental cell lysis time statistics of *λ*-phage mutants^40^. In the presence of post-transcriptional regulations by micro-RNA^49^, again for bursty protein production, relevant FPT distributions have been theoretically studied^50^.

The above being the status for proteins, for the far simpler case of threshold crossing of mRNA copy number, where transcription involves just a single-stage production and degradation, the full FPT distribution is not explicitly known – we provide this exact result in this paper. We note that a few moments of this FPT problem are well known^51^, while computational and approximate analytical studies of the first passage of mRNA were done, even with more complicated promoter kinetics^52–54^. As various technologies of monitoring mRNA levels in cells exist, the FPT distribution may be of practical use. Large fluctuations in mRNA distributions, whether in copy number or FPT, are directly tied to protein translation. In fact, a very short time threshold crossing of mRNA may trigger quicker post-transcriptional regulations by micro-RNA or P-bodies to regulate protein translation.

The paper is organized as follows. In Sec. II, we introduce the transcription model, and define the mRNA threshold crossing as a first passage problem. In Sec. III we present the FPT distribution in Laplace space following a backward time formalism. In Sec. IV we compute the relative fluctuation measures and show that they have *U*-shapes. In Sec. V we extend the exact formula for FPT to include the DNA duplication mechanism, and corresponding doubling of transcription rate. In Sec. VI we summarize our results.

## II. THE MODEL FOR TRANSCRIPTION AND THE M-RNA THRESHOLD CROSSING PROBLEM

The model we study here involves the transcription of mRNAs at a rate *k* and their degradation at a rate *γ*. In the absence of any promoter regulation by transcription factors or nucleosome binding-unbinding, this is the standard mRNA synthesis model. Even for this simple model, which has been widely studied in the gene expression literature^46,55^, while theoretically finding the distribution of mRNA copy numbers is easy, the threshold crossing time distribution is nontrivial as we will see below. A schematic diagram of the model is shown in Fig.1(a).

**FIG. 1:**
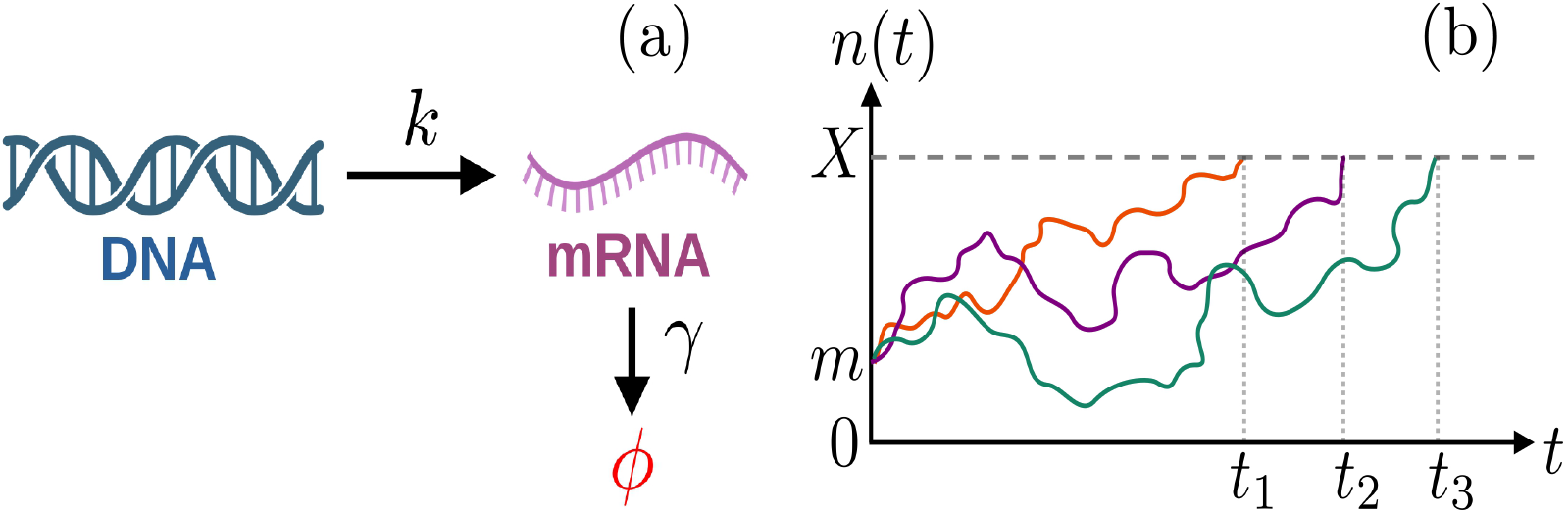
(a) A schematic diagram of the model of gene transcription with transcription rate *k* and degradation rate *γ* per mRNA molecule. (b) A schematic diagram of the first passage process, showing three representative trajectories of mRNA count *n*(*t*) versus time *t* starting from *m* to reach the threshold *X* – the corresponding times of process termination are *t*_1_, *t*_2_ and *t*_3_.

Let, at any time *t*^′^, *n* be the number of mRNA copies of a gene present in the cell, given *m* mRNAs at an initial time *t* – the associated probability is *P*(*n, t*^′^|*m, t*). Some representative trajectories and the corresponding FPT are shown schematically in Fig.1(b). The governing forward Master equation^51,55^ predicting future evolution of *n* following this probability distribution is given by

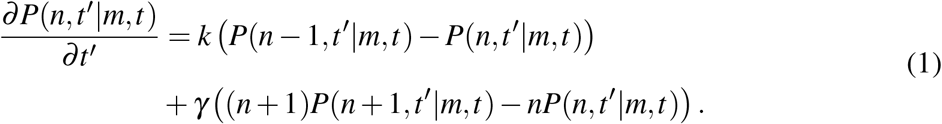

Solving this equation with boundaries *n, m* ∈ [0, ∞) is trivial, and particularly for *m* = 0 it gives the well known Poisson distribution for mRNA copy number^51,55^.

However, one is often interested in the first passage problem and the distribution of first passage times (FPT) in which the mRNA count *n* crosses a fixed threshold *X* for the first time. Thus, the boundaries are now *n, m* ∈ [0, *X*]. Without any loss of generality, we set *t*^′^ = 0 and *t* → −*t*, and are left to estimate the unknown statistics of the FPT *t* until which the survival continues, i.e., the threshold crossing does not happen. One defines the survival probability as 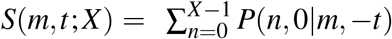 and at the boundary *S*(*m* = *X, t*; *X*) = 0 ∀ *t* along with the initial condition *S*(*m*, 0; *X*) = 1 ∀ *m* ∈ [0, *X* − 1]. The FPT distribution is related to the survival probability as *f* (*m, t*; *X*) = −*∂S*(*m, t*; *X*)/*∂t*. Since we are concerned with survival from the past for first passage times, the backward Master equation evolving *P*(*n, t*^′^|*m*, −*t*) in time *t* leads to the equation for survival probability^51^:

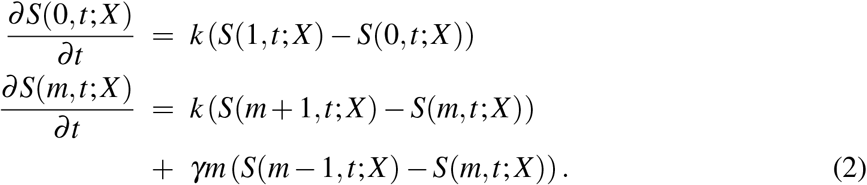

The second equation above is valid for all the integers *m* ∈ [1, *X* − 1], and for *m* = *X* − 1, *S*(*m* + 1, *t*; *X*) = 0 (absorbing boundary). From (2), one can obtain the difference equations for mean FPT 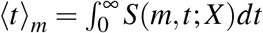, and the second moment 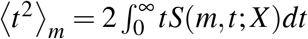 as follows:

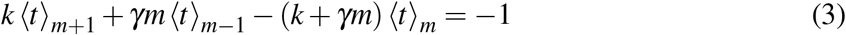

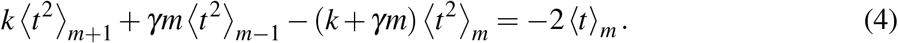

The absorbing boundary conditions, ⟨*t*⟩_*X*_ = 0 and ⟨*t*^2^⟩_*X*_ = 0 may be used to solve those – see the next section for their explicit expressions.

The point, however, in this paper is not to stay restricted to the first two moments, but to obtain the full statistical information encoded in the distribution function of FPT.

In the next section, we follow the backward time formalism, which directly provides the survival probability and FPT distribution, not via *P*(*n, t*^′^|*m, t*) as in the forward time formalism. In principle, the forward formalism is also a valid approach, and using Eq. 1 one may formally write the time dependent distribution – see Appendix A. However, to find the explicit FPT distribution in real time *t*, the challenge is to find the eigenvalues *λ*_*i*_ and eigenvectors 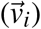 of the relevant W-matrix. Given that, for a tridiagonal matrix with unequal coefficients, as in this case, this is not generally easy, it remains an open problem in the literature.

In contrast, as we show below, in the Laplace space the challenge is not of a matrix diagonalization but of matrix inversion, which is a lot easier. We find it particularly straightforward to do so for the backward time equations for the survival probabilities, starting with Eq. 2. It is also possible to arrive at the same answer through an alternative forward time formalism but in Laplace space, as we show in Appendix A. We note that the information content of the FPT distribution remains the same whether the answer is in real time or in the Laplace domain.

For completeness, we briefly note that for mRNA ‘concentration’ *c*(*t*), rather than the Mrna number, crossing a threshold *c* = *X*, the following is the forward Fokker-Planck equation, which is the continuum analog of Eq. 1:

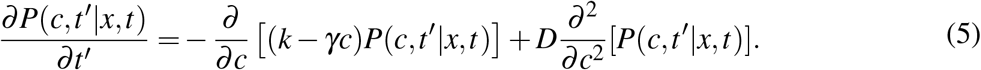

For the case of *k* = 0, the problem reduces to the classic Orstein–Uhlenbeck process and has been analyzed in^56,57^. Yet for *k*≠ 0, the FPT distribution in only known in Laplace space from a recent work^40^ and is:

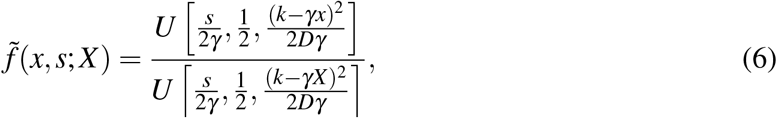

where *U* is the Tricomi function^58^.

## III. THE THRESHOLD CROSSING TIME DISTRIBUTION IN LAPLACE SPACE

We define the Laplace transforms of the survival probabilities as 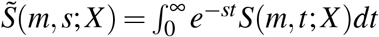 and of the FPT distributions as 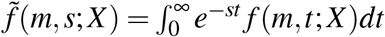. Taking Laplace transform of Eq.2 and using 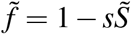 (since *f* = −*∂S*/*∂t*), we obtain

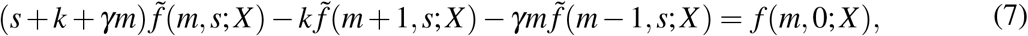

where, *f* (*m*, 0; *X*) = 0 for *m*≠ *X* . Starting at the absorbing state, first passage occurs immediately, hence *f* (*X, t*; *X*) = *δ* (*t*) leading to 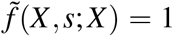. For the transient states *m* = 0, …, *X* − 1, the above set of equations can be written as follows involving a matrix A:

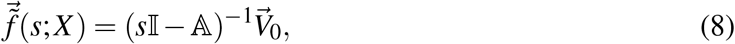

where vectors 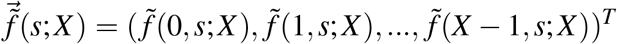 and 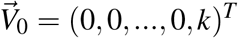. This particular form of 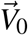 which follows from the absorbing boundary condition, implies that only the last column elements of (*s*I −A)^−1^ are required to calculate the full FPT distribution, i.e.,

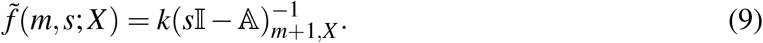

The matrix sI−A is of the tridiagonal form,

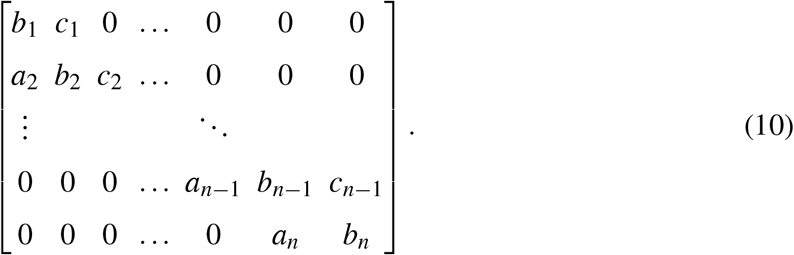

where the specific elements are *a*_*i*_ = −(*i* − 1)*γ, b*_*i*_ = *k* + *s* + (*i* − 1)*γ* and *c*_*i*_ = −*k*. The inverse of a tridiagonal matrix is explicitly known from the literature and can be expressed in terms of the determinants of the leading(*U*_*i*_) and trailing (*L*_*i*_) principal sub-matrices^59,60^. In Appendix B, the particular element 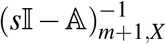 (in Eq.9) is evaluated and that yields:

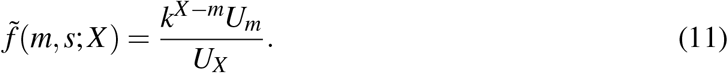

To proceed further, one needs the explicit form of the determinants *U*_*m*_, which satisfy a recurrence relation

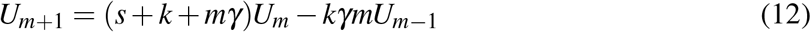

with *U*_0_ = 1, *U*_1_ = *s* + *k* and *U*_2_ = *k*^2^ + 2*ks* + *s*^2^ + *sγ*. We recognize the pattern of *U*_*m*_ and propose a general formula:

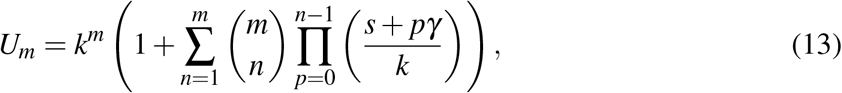

which we then prove by mathematical induction based on Eq. 12 in Appendix B. The above formula appeared in a queuing theory problem which was proved by a different method^61^. Substituting the expression for *U*_*m*_ from Eq. 13 in Eq.11 gives the main result of our interest, namely the exact FPT distribution in Laplace space:

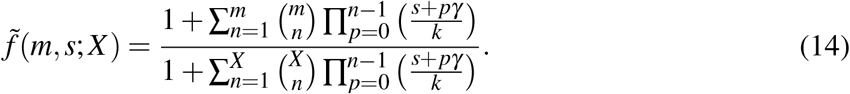

The time dependent FPT distribution *F*(*m, t*; *X*) may be obtained by numerical inversion of the above Laplace transform 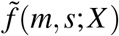, using **Mathematica**^62^.

Fig.2 shows the distributions *F*(*m, t*; *X*) of FPT times *t*, for two threshold values *X* = 35 and *X* = 70 with initial *m* = 10 mRNAs. For *X* = 35, the distribution is relatively narrow and sharply peaked, indicating comparatively small fluctuations in the first passage time. In contrast, increasing the threshold to *X* = 70 results in a broader and more skewed distribution, indicating large fluctuations in time required to cross the threshold. We have also performed Gillespie simulations^63^ for the mRNA transcription model, and obtained the threshold crossing times – the normalized histograms thus obtained (in gray and blue colors) match perfectly the two analytically predicted curves (solid lines in Fig.2). We now proceed to study the measures of relative fluctuation which follow from the exact formula Eq. 14.

**FIG. 2:**
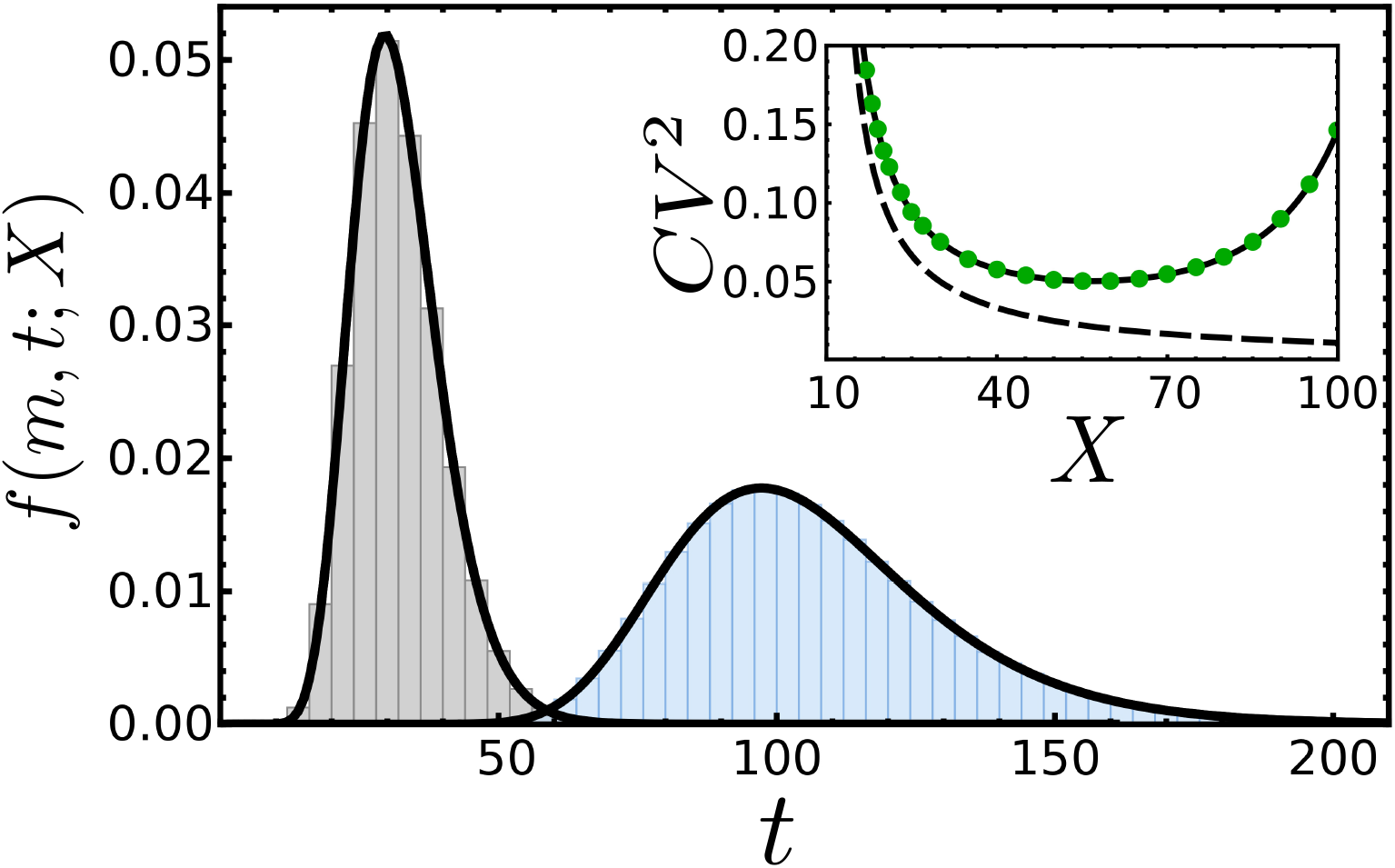
FPT distributions for two different thresholds *X* = 35 (left) and *X* = 70 (right). The initial count of mRNA is *m* = 10 in both the cases, and rates *k* = 1.0 min^−1^ and *γ* = 0.01 min^−1^. The solid black lines show the analytical expression (Eq. 14), while the Gillespie simulation data is shown as normalised histograms in Gray (left) and blue (right). Inset: Variation of *CV*^2^ with threshold *X* showing a non-monotonic behavior – theoretical curve (using Eq. 16,17) is in solid line and Gillespie data in green symbols. The dashed line is for *γ* → 0, showing a monotonic decrease.

## IV. THE MEASURES OF RELATIVE FLUCTUATION OF THRESHOLD CROSSING TIMES AND THEIR U-SHAPES

While the FPT distribution has full statistical information, in experiments, often partial information in the form of moments and cumulants is of interest, as they show interesting trends as a function of tunable parameters. For example, the first passage times of cell lysis in *λ*-phage infected E. coli due to holin protein concentration crossing a threshold, the coefficient of variation *CV*^2^, skewness, and decay times of exponential tails with respect to mean (*τ*/⟨*t*⟩) – all showed non-monotonic behavior as a function of the threshold concentration *X* ^38,40^. Similarly, it was theoretically shown that in the presence of post-transcriptional regulation by micro-RNA, the threshold crossing times of bursty proteins show a non-monotonic *U*-shape for *CV*^2^, skewness and *τ*/⟨*t*⟩ as a function of the rates of binding and unbinding of the microRNA-mRNA complex^50^. The mRNA threshold crossing problem at hand is less complex compared to the above problems, and thus, it is interesting to study the measures of relative fluctuations to see if similar trends exist here in this simpler case.

We note that the moment of any order, namely the p-th, can be obtained from the exact (Laplace) FPT distribution Eq.14 by taking derivatives:

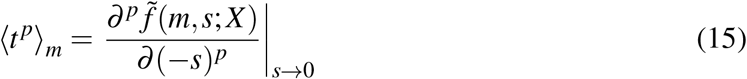

For notational brevity we define *D*_*m*_(*s*) = *U*_*m*_(*s*)/*k*^*m*^ where *U*_*m*_ follows from Eq. 13. Accordingly the coefficients 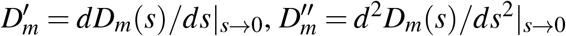 and 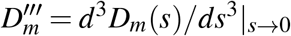. Using these, the expressions of mean, variance, and third cumulant are

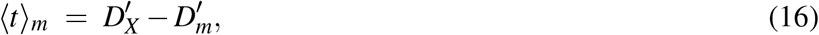

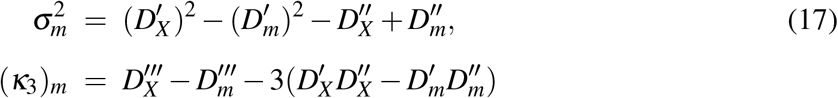

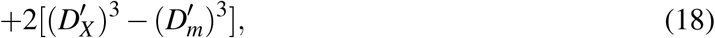

where the required coefficients are

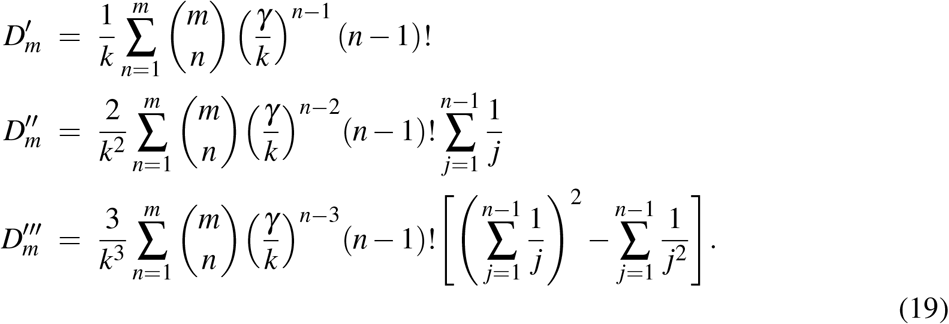

In the inset of Fig. 2 we show the plot of analytical 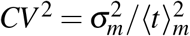 versus *X* (using Eqs. 16,17) in solid line, along with Gillespie simulation data (in green symbols) supporting it. The *U*-shape is to be noted, which is reminiscent of similar shape found for *CV*^2^ of protein threshold crossing times in earlier literature^37–40,50^. As in case of proteins, the *U*-shape owes its origin to the degradation factor *γ* – as *γ* → 0, the inset of Fig. 2 shows that the curve (dashed line) becomes monotonic. At large *X* the number of stochastic steps needed to cross the threshold is typically large with correspondingly high mean FPT. Over such prolonged duration degradation (for *γ*≠ 0) causes occasional drift away from the target threshold — this leads to high trajectory to trajectory variability in time, leading to the U-turn upwards in the *CV*^2^ curve^50^.

Likewise, the skewness of the mRNA threshold crossing 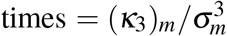 also shows an *U* shape (Fig.3(a)), just as in case of proteins^40,50^. The solid line in the figure follows from Eq. 18, while the symbols (in red) are from Gillespie simulations.

**FIG. 3:**
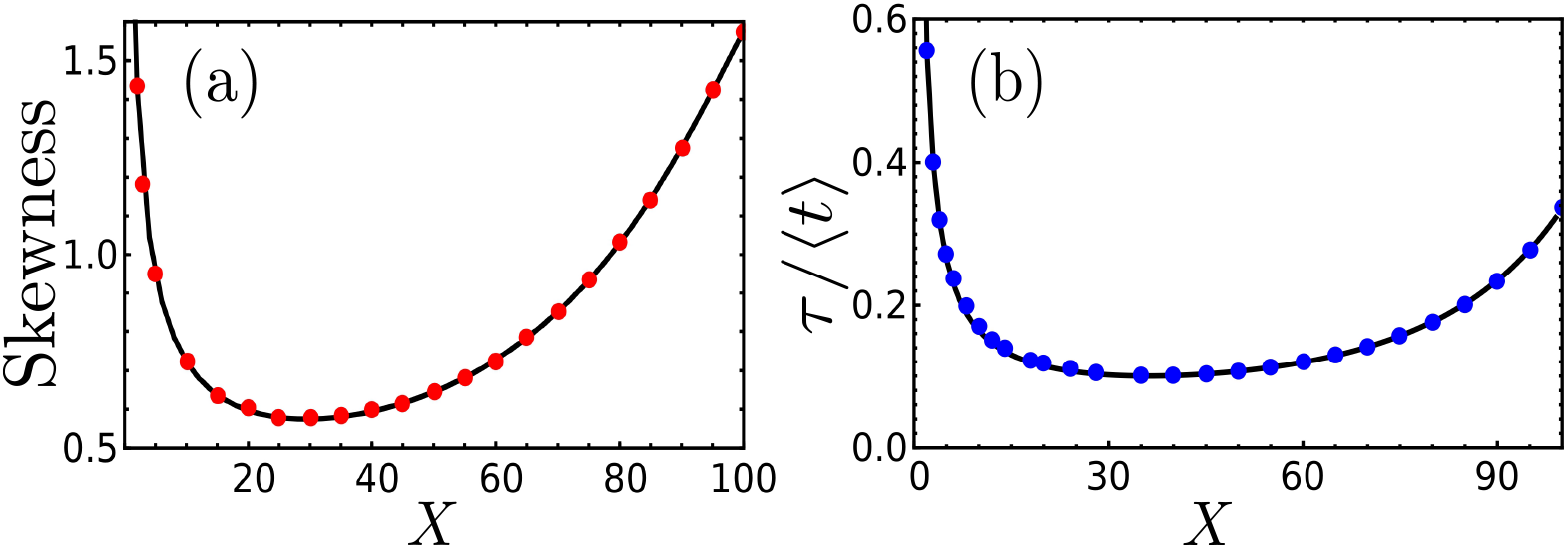
Variation of (a) skewness and (b) relative characteristic time (*τ*/⟨*t*⟩) with the threshold number *X* . The rates *k* = 1.0 min^−1^, *γ* = 0.01 min^−1^, and initial *m* = 0. The black solid lines are obtained from analytical expressions and colored circles are Gillespie simulation data.

Finally, we note that the FPT distribution *f* (*X, t*; *m*) has an exponential tail ∼ exp(−*t*/*τ*_*c*_), where *τ*_*c*_ is referred to as the characteristic time^34,50^. The latter is determined by the pole in the denominator of Eq.14 whose real part, −*α*_*c*_, has the lowest absolute magnitude:

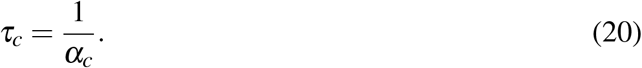

Note that finding the roots of the denominator in Eq.14 analytically remains difficult – in fact, if it were possible for arbitrary *X*, the time dependent explicit form of *f* (*X, t*; *m*) would have been immediately obtained by the method of residues. So we obtain *α*_*c*_ numerically and plot it as a solid line in Fig.3(b); the corresponding Gillespie data are shown in blue symbols. We observe again a U-shape, reminiscent of the characteristic FPT of proteins^40,50^.

## V. DNA DUPLICATION AND MRNA FPT DISTRIBUTION

So far we considered mRNA transcription at a constant rate *k* until the threshold *n*(*t*) = *X* is reached. However, as eukaryotic cells make a transition from *G*1 phase to *S* phase in their cell cycle, the DNA duplicates and as a result the rate of mRNA transcription doubles (*k* → 2*k*) assuming sufficient RNA-polymerase is present^64^. In prokaryotic cells too a similar thing happens as the cell transitions from *B* to *C* phase^65^. Suppose the DNA duplication happens at time *t*_0_ and the corresponding mRNA copy number is *n*(*t*_0_) = *N*. The FPT distributions derived in this paper needs revision if the target threshold *X* > *N*, as the transcription rate changes for the process as *n*(*t*) grows from *N* to *X* . Note that the time *t*_0_ at which duplication happens is highly variable due to many factors^66^, and accordingly is the corresponding *N* (which is also different for different genes within the cell).

Studies on variability of gene products have incorporated the effect of DNA duplication in multi-stage models of gene expression^11^. Here we are concerned with first passage process, where time is not fixed to find a distribution of *n*, but rather a threshold is fixed to find the distribution of times of passage. In that spirit, we propose a way to extend our formalism to study gene duplication. We assume that for a concerned gene, there is a given *N* at which the duplication happens – although biologically there may be a distribution of *N*, for simplicity we ignore that. Then, if *t*_1_ is the time taken for mRNA count to react *N* starting at 0 for the first time (at transcription rate *k*), and *t*_2_ is the time to reach *X* starting at *N* (at transcription rate 2*k*), the total FPT *t* = *t*_1_ + *t*_2_. Hence the desired FPT distribution, which we denote by *F*(*m, t*; *N, X*), is a convolution of two FPT distributions *f*_*k*_(*m, t*_1_; *N*) and *f*_2*k*_(*N, t*_2_; *X*), and is given by:

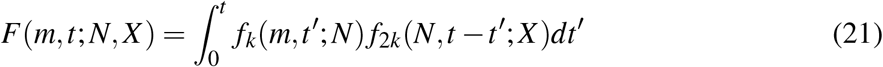

Hence the Laplace transform 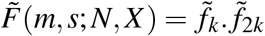 is a product. Here the subscripts (*k* and 2*k*) to the *f* functions remind of the different transcription rates involved in the two stages, pre- and post-duplication. The Laplace transform 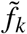 follow the expression in Eq. 14, and 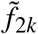 does the same with replacement of *k* → 2*k*. Thus we have,

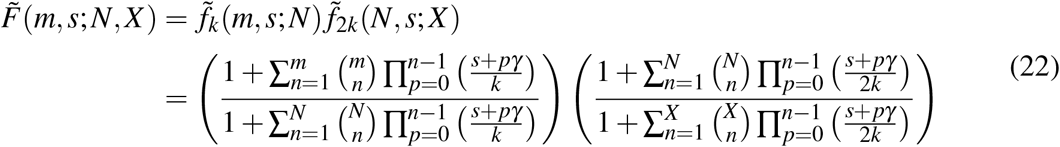

Note that on an average, *N* is expected to have a monotonically increasing relationship with *t*_0_. Hence by varying *N*, we may study how an early or a late duplication affects the FPT distribution for a fixed threshold *X* . In Fig.4 we present the inverted FPT distrbution *F*(*m, t*; *N, X*) using the exact expression Eq. 22 for *N* = 20 and 70 (given *X* = 90); normalised histograms from Gillespie simulations (in yellow and purple colors) perfectly match the theoretical curves. For *N* = 20 as the second stage with rate 2*k* dominates, the distribution has a smaller mean and visibly small variance, compared to the *N* = 70 case where the first stage with rate *k* dominates. Interestingly we find in the inset of Fig.4 that as a function of *N*, the *CV*^2^ of *F*(*m* = 0, *t*; *N, X*) has an *U*-shape.

**FIG. 4:**
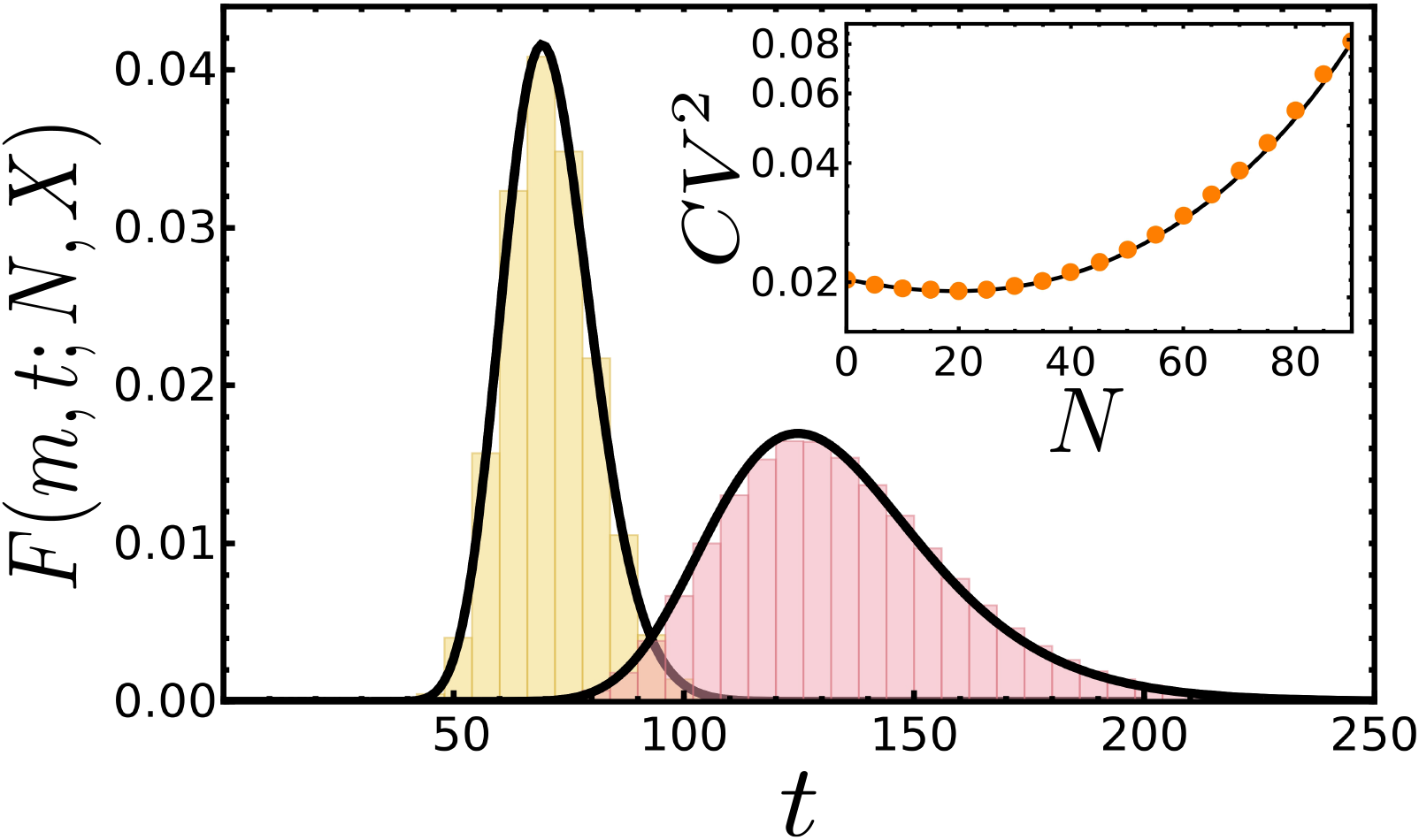
FPT distribution *F*(*m* = 0, *t*; *N, X*) versus time *t* for *N* = 20 (left) and *N* = 70 (right). The threshold *X* = 90, and rates *k* = 1.0 min^−1^, *γ* = 0.01 min^−1^. The solid lines are theory (see text) while, histograms are Gillespie simulation data. Inset: *CV*^2^ of the *F*(*m* = 0, *t*; *N, X*) distribution versus *N* shows an *U*-shape (for the same parameters as in the main figure).

## VI. SUMMARY

The simple model of transcription with constitutive mRNA production and degradation is well known and easily solved problem for its copy number distribution. Yet the corresponding threshold crossing problem is challenging, and we have provided an exact analytical answer for the Laplace transform of the first passage time distribution. We have also extended the distribution by incorporating the effect of transcription rate switching due to DNA duplication. The results add to the developing literature on first passage of protein levels. Although protein threshold level crossings are known to directly trigger critical biological events, mRNA levels are also important for post-transcriptional regulation by micro-RNAs and P-bodies which affect subsequent protein translation. Thus knowledge of a distribution of level crossing times of mRNA is expected to have practical use.

The explicit FPT distributions in Laplace domain (Eq.14 and Eq. 22) contain all the statistical information. The time-dependent distribution is readily obtained by inversion using **Mathematica**^62^. Specifically the Laplace transforms enables one to obtain moments of arbitrary order, thereby providing mean, variance, and higher cumulants of the FPT, as presented in this paper. Moreover large deviations are captured through the characteristic time of the tail of the FPT distribution and we have shown how that may be obtained from the the poles of the Laplace FPT distribution. At every step, for different distributions and cumulants, we have provided supporting Gillespie simulation results, to confirm our analytical predictions.

An interesting observation is the existence of non-monotonic behaviour of relative fluctuations of the FPT. Particularly, *CV*^2^, skewness and the characteristic time relative to mean time all exhibited non-monotonic U-shapes. This as we have shown is a consequence of the non-vanishing degradation factor, which enhances fluctuations at high thresholds. It is also significant that similar behavior have been reported for more complicated models of threshold crossing in proteins.

We incorporated the switching of mRNA transcription rate due to DNA duplication within a cell cycle. We argued that this switch happens when mRNA number crosses an intermediate threshold number (lower than the final threshold number). We observed that an early or intermediate duplication results in dynamics dominated by the enhanced transcription rate 2*k*, yielding a quicker passage and small fluctuations. Whereas late duplication results in dominance of a smaller transcription rate *k*, leading to a delayed passage and greater fluctuations. There is a *U*-shape of the associated *CV*^2^ as a function of the intermediate threshold number.

We hope that the exact FPT distributions of mRNA would be useful for theorists and experimentalists in the field of stochastic gene expression.

## ACKNOWLEDGMENTS

AKV and HKB acknowledge fellowship support from the University Grants Commission (UGC), India, Ref. numbers 221610137247 and 211610033150, respectively, and support from IIT Bombay.

## AUTHOR DECLARATIONS

### Author Contributions

**Arunendra Kumar Verma**: Conceptualization(equal); Data Curation(equal); Formal Analysis(lead); Visualization(equal); Writing/Original Draft Preparation(equal); Writing/Review & Editing(equal) **Hillol Kumar Barman**: Conceptualization(equal); Data Curation(equal); Formal Analysis(equal); Software(lead); Visualization(equal); Writing/Original Draft Preparation(equal); Writing/Review & Editing(equal) **Krishna Rijal**: Conceptualization(equal); Data Curation(equal); Formal Analysis(equal); Visualization(equal); Writing/Original Draft Preparation(equal); Writing/Review & Editing(equal) **Dibyendu Das**: Conceptualization(equal); Data Curation(equal); Formal Analysis(equal); Supervision(lead); Visualization(equal); Writing/Original Draft Preparation(equal); Writing/Review & Editing(equal)

## DATA AVAILABILITY

The data that support the findings of this study are available from the corresponding author upon reasonable request.

## Appendix A: Forward time formalism and FPT distribution

The Eq. 1 may be cast into a form involving a vector and a matrix as follows:

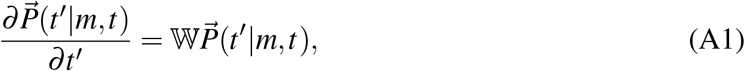

where 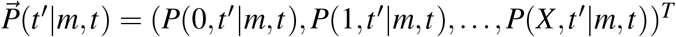 . The inclusion of the last component *P*(*X, t*^′^|*m, t*) in the vector 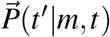 saves the trouble of putting in the boundary condition, and at long time, all probability flows to the *n* = *X* state. In particular,

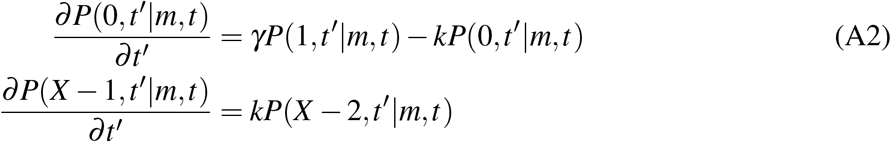

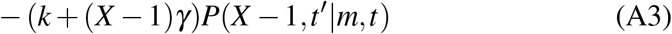

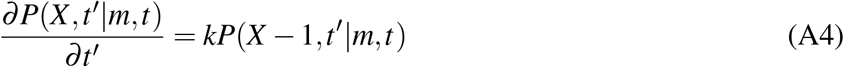

The initial condition is *P*(*n, t*|*m, t*) = *δ*_*n,m*_. The formal solution to Eq. A1 is given by 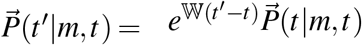 . The explicit form of the matrix W is

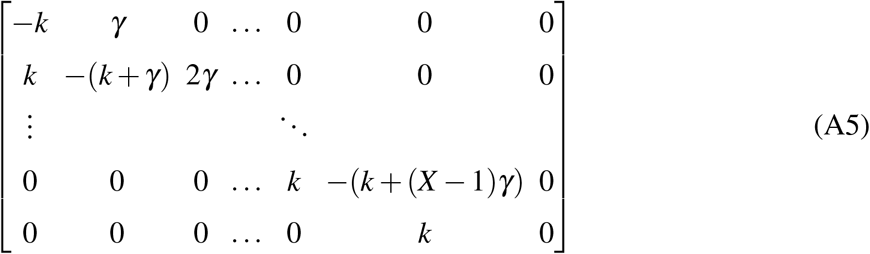

which is a tridiagonal matrix of dimension (*X* + 1). It is apparent that one of the eigenvalues is 0 and the corresponding eigenvector *v*_0_ = (0, 0, 0,, 1)^*T*^, which is the steady state corresponding to eventual absorption of mRNAs at the threshold boundary *X* . The remaining eigenvalues are *λ*_*i*_ < 0 and they have corresponding eigenvectors 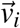 .

A formal solution can be expressed in terms of the eigevectors 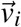,

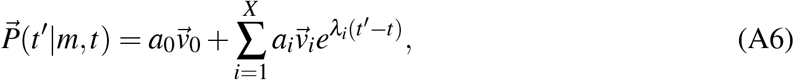

where the constants *a*_*i*_ are determined from the initial condition. But the challenge of this approach is knowing the eigenvalues and eigenvectors of W, which is not straightforward for this tridiagonal matrix with unequal elements. This makes it hard to obtain the FPT distribution in real time domain.

An alternative is to work in the Laplace domain, and we choose *t* = 0. Taking the Laplace transform of the Eq. A1, with 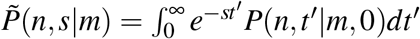 and defining vectors 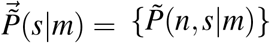 and 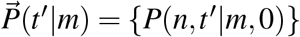 gives

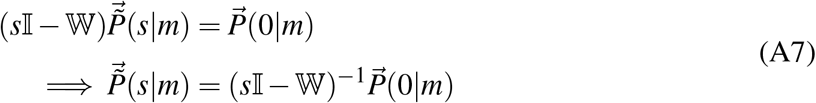

The matrix *s*I −W retains the tridiagonal form and is explicilty given by,

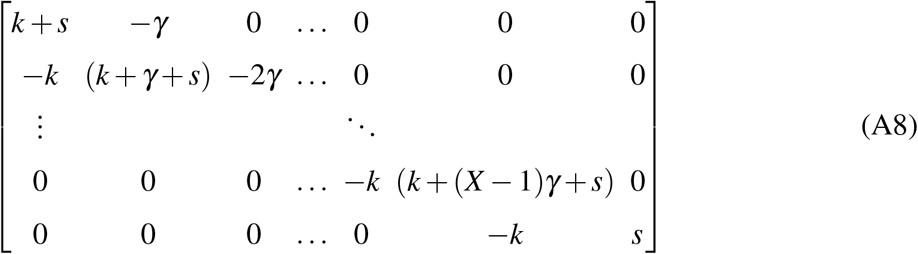

The initial probability vector 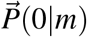 contains only one non-zero element namely 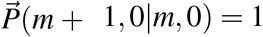, hence the inverse selects only the (*m* + 1)th column and one may write,

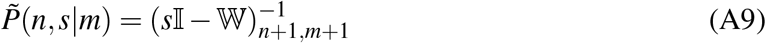

Particularly, the survival probability *S*(*m, t*; *X*) = 1 − *P*(*X, t*^′^ = *t*|*m*, 0), and FPT distribution *f* (*m, t*; *X*) = *kP*(*X* − 1, *t*^′^ = *t*|*m*, 0). Hence, we obtain the FPT distribution in Laplace space as follows:

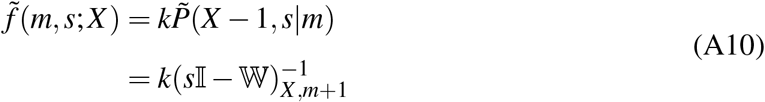

One observes that for the transient states, i.e. *n* ∈ [0, *X* − 1], the form of the matrix *s*I − W is simply the transpose of *s*I − A, the matrix (Eq. 10) we studied within the backward formalism. This provides a convenient comparison to show the equivalence of the two methods. Since the determinant is unchanged during a transpose, the determinant of the leading principal submatrices 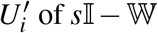 are the same as the corresponding *U*_*i*_ of *s*I −A, in particular

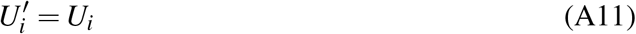

and the determinant of the full *X* + 1 dimensional matrix *s*I −W is

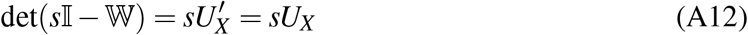

Using the general expression for the inverse of a tridiagonal matrix, discussed in Appendix B, we see that the first passage probability distribution can be written as,

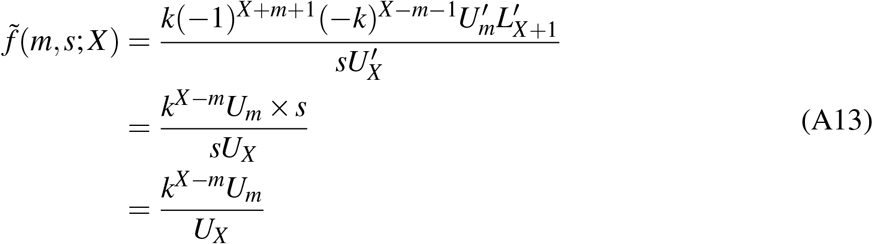

where 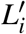 are the trailing principal submatrices of *s*I − W and in particular 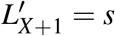. Hence, the result so obtained is identical to the result obtained using backward formalism (see Eq. 11). We now proceed to prove the explicit *U*_*m*_ form in Eq. 13.

## Appendix B: General form of *U*_*i*_ and the FPT distribution

As discussed in the main text, the matrix *s*I−A, is a tridiagonal matrix, and the FPT distribution in the Laplace domain is obtained from a particular element of (*s*I −A)^−1^.

The inverse of a general tridiagonal matrix can be expressed in terms of the determinants of its leading and trailing principal submatrices^59,60^. For a general tridiagonal matrix, B, of dimension *Q*, with diagonal elements *b*_*i*_, subdiagonal elements *a*_*i*_ and superdiagonal elements *c*_*i*_, as shown in Eq.10, the inverse elements are given by,

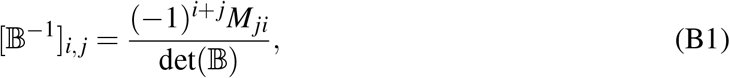

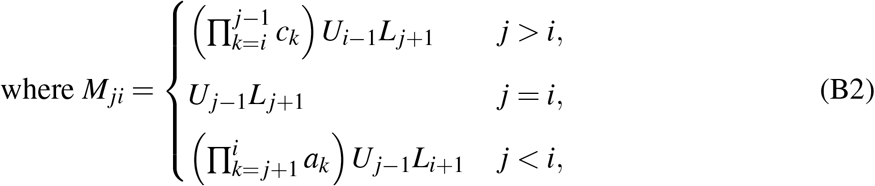

Here, *U*_*i*_ and *L*_*i*_ denote the determinants of the leading and trailing principal submatrices, and follow the following recurrence relation for a *Q* dimensional matrix,

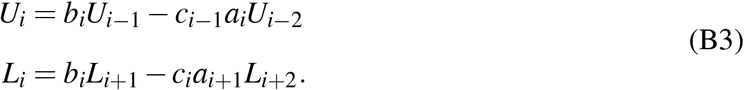

In particular, *L*_*Q*+1_ = 1, *L*_*Q*_ = *b*_*Q*_, *U*_0_ = 1, *U*_1_ = *b*_1_ and *U*_*X*_ = det(B). For the matrix *s*I − A, we have *a*_*i*_ = −(*i* − 1)*γ, b*_*i*_ = *k* + *s* + (*i* − 1)*γ* and *c*_*i*_ = −*k*, and the explicit structure of the said matrix is

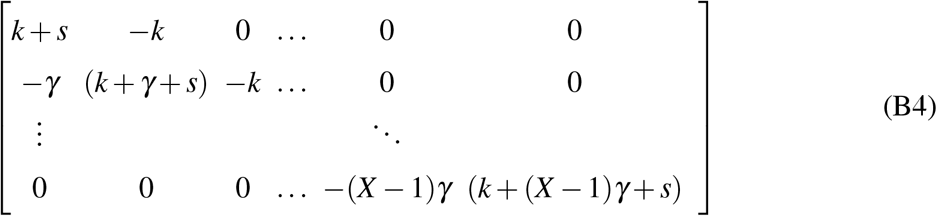

Thus, *s*I − A is a tridiagonal matrix of dimension *X* and we require its inverse to obtain the distribution. From Eq.9 we see that one only requires the last column of the inverse matrix (*s*I − A)^−1^, and from Eqs.B1, B2 one gets 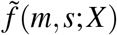 to be:

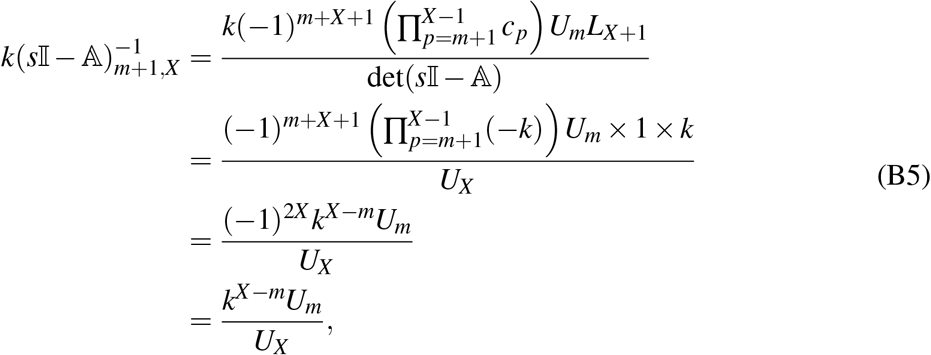

same as in Eq. 11. Now we need the explicit form of the coefficients *U*_*m*_, which is the main aim of this Appendix. The first few coefficients are:

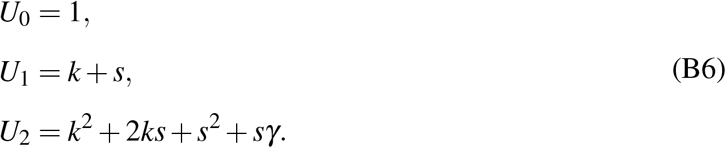

The above equations show that our proposed general form of *U*_*m*_ in Eq. 13 is satisfied for *m* = 1 and *m* = 2. We now proceed to prove it by mathematical induction. We assume that it holds true for *m* = *N* − 1 and *m* = *N* and show that it holds for *m* = *N* + 1. The recurrence relation for *U*_*m*_ (Eq.12) imply

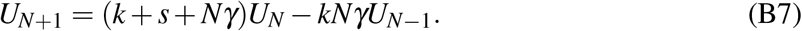

Substituting the terms *U*_*N*−1_ and *U*_*N*_ from the induction hypothesis we see,

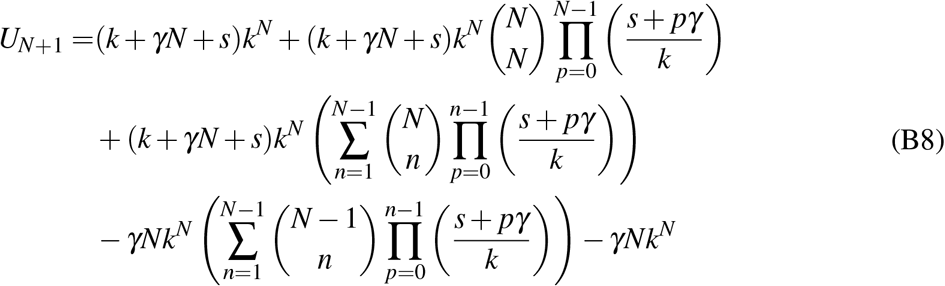

Combining the third and the fourth term and invoking the identity 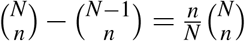, we obtain

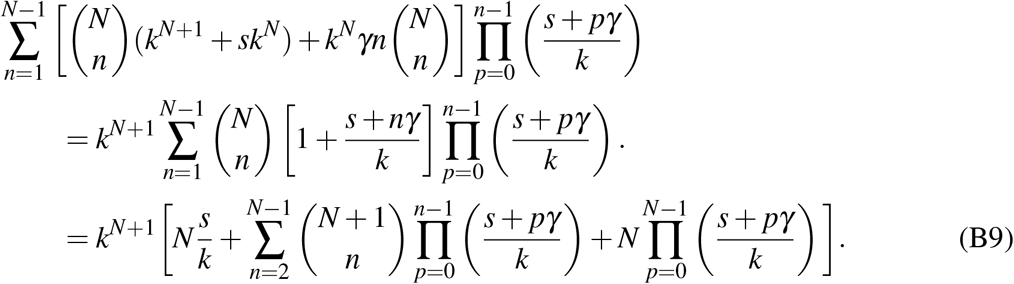

Combining this term with the remaining terms yields,

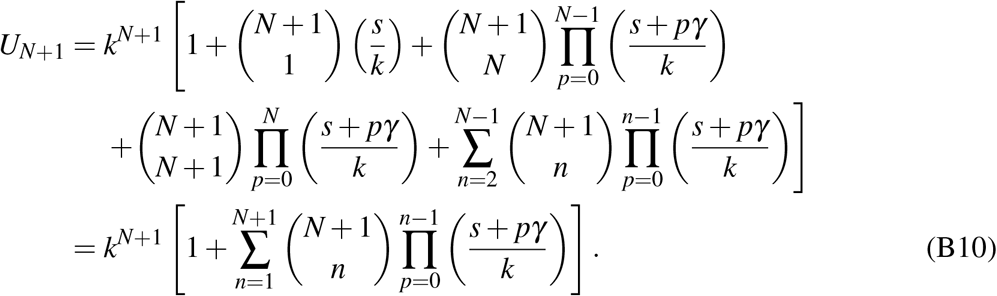

Thus we see that Eq.13 holds for *U*_*N*+1_ proving that the proposed form for *U*_*m*_ is true. As a consequence 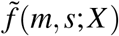 is given by Eq. 14.

